# Glioblastoma exhibits bidirectional adaptation to treatment through JARID2-associated developmental plasticity

**DOI:** 10.64898/2026.11.17.766938

**Authors:** Morgan Thomas, Bo Wang, Isabel Craghill, Yousef Alghamdi, Muna Al-Jabri, Udayaraja GK, Martina Finetti, Steven Pollock, Michael D Cearns, Laura Matthews, Thomas A Hughes, Michael D Jenkinson, Andrew R Brodbelt, Mark Wade, Abhinav K Jain, Andy Bulpitt, James Poulter, Lucy F Stead

## Abstract

Glioblastoma (GBM) exhibits bidirectional transcriptional reprogramming following standard-of-care (SoC) treatment, yet the direction cannot be predicted from primary tumours. Using independent longitudinal bulk and single-cell datasets, we show that this response is captured by a framework, the JARID2 Binding Site (JBS) axis. Tumours and cells can be positioned along this axis according to expression of JBS genes, with responder subtypes defined by movement along the axis from primary to recurrent states. The JBS axis aligns with established GBM ecosystems and their constituent cell populations, and recurrent tumours polarise towards opposite ends. Changes in tumour microenvironment composition strongly influence this polarisation and drive adaptive transcriptional responses more profoundly than chemoradiation alone. Mechanistically, JBS genes are enriched for promoters marked by persistent bivalent chromatin and neurodevelopmental regulatory programmes. These findings suggest that JARID2-associated Polycomb regulation maintains a reservoir of developmental plasticity that enables treatment-driven ecosystem adaptation and represents a potential therapeutic vulnerability.

**Statement of significance:** We propose that JARID2-associated promoter bivalency constitutes an epigenetic substrate for ecosystem remodelling in glioblastoma, enabling diverse recurrence trajectories but via a conserved and potentially targetable underlying mechanism of adaptation.

**Conflicts of Interest:** The authors declare no potential conflicts of interest

## Introduction

Glioblastoma (GBM) remains an incurable brain tumour, with most patients experiencing a subsequently fatal recurrence within 7 months of receiving standard of care (SoC) treatment of maximal safe surgical resection and subsequent chemoradiotherapy(1). Multiple longitudinal genomic, transcriptomic and single-cell studies have sought to understand how unresected GBM cells evolve and adapt to survive adjuvant treatment, suggesting that this is not driven by clonal selection alone(2–4). Instead, coordinated changes in transcriptionally-defined malignant cell states and tumour microenvironment composition supports a model in which recurrence is driven by ecosystem remodelling(5–10). Collectively these findings suggest that GBM recurrence does not result from convergence upon a common terminal phenotype but is facilitated by continual adaptive state transition. The mechanisms that permit such ongoing conversion between cellular states and ecosystem configurations remain poorly understood.

Whilst DNA methylation profiles remain largely stable through SoC treatment, the observed changes in neoplastic cell state in the absence of consistent genetic alterations implicates alternative epigenetic mechanisms underpinning treatment resistance(11,12). It has recently been suggested that cytotoxic therapy induces persistent chromatin alterations capable of reshaping GBM cell transcriptional programmes long after treatment exposure(13). This raises the possibility that recurrence-associated ecosystem remodelling may be mediated by semi-stable epigenetic mechanisms that are influenced by exposure to chemoradiation. We previously identified a reproducible transcriptional response to SoC treatment in GBM involving genes containing JARID2 binding sites (JBS genes)(7). JARID2 is a key player in chromatin remodelling during developmental gene regulation via interaction with Polycomb Repressive Complex 2 (PRC2)(14,15). We showed that JBS gene dysregulation stratified GBM patients according to whether transcriptional reprogramming post-SoC treatment, comprised up- or down-regulation, rather than the absolute molecular state of their tumour at either timepoint(7).

Here we expand upon our previous work to show that JBS gene expression, and changes therein, frame a continuous adaptive axis spanning proneural and mesenchymal ecosystem states in GBM. By integrating bulk RNA sequencing, single-cell datasets, spatial transcriptomics and longitudinal chromatin profiling, we demonstrate that ecosystem remodelling occurs along this axis and is associated with persistent bivalent chromatin states at Polycomb-regulated developmental loci. In this way, JBS genes do not define a specific GBM state, they define a recurrence-enabling plasticity programme, which presents a novel therapeutic vulnerability.

## Results

### GBM tumours consistently transcriptionally reprogramme in response to SoC

We previously identified a subset of genes that are consistently dysregulated between paired primary and recurrent GBM tumours(7). These all contain JARID2 binding sites in their promoters, so were coined JBS genes. Based on the direction of change, patients stratify into Up responders (JBS genes increase at recurrence) or Down responders (JBS genes decrease at recurrence). Additional independent bulk and single cell longitudinal GBM transcriptional datasets have been created since we first described the responder subtype phenomenon(7–10). We acquired and analysed these new datasets, which confirmed the presence, and proportion, of Up and Down responders as per our original finding (Fig.1A, Supp.Table.1). Having shown that responder subtypes appear to adopt distinct mechanisms of treatment resistance (Fig.1B)(7), we proceeded to investigate whether GBMs can be classified at primary diagnosis to inform personalised medicine strategies

**Figure 1.**
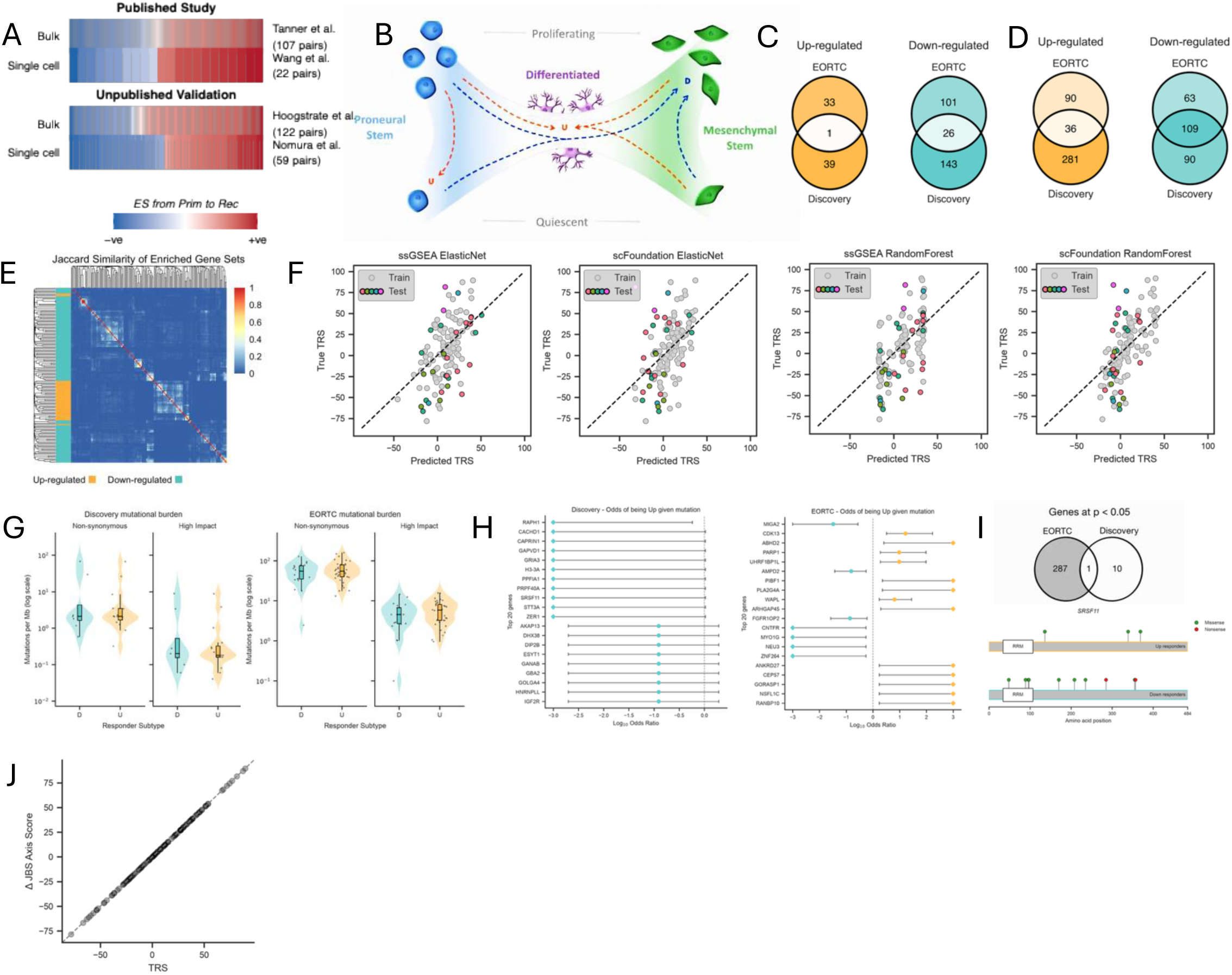
A. The responder subtype phenomenon is validated in both bulk and single cell longitudinal GBM datasets published since our initial discovery of it [ES; Enrichment Score]. B. Phenotypic adaptions observed in responder subtypes map to bona fide resistance mechanisms (taken from Tanner et al, 2024). C. Overlap in genes significantly rising or falling with TRS in primary tumours in two independent cohorts (Discovery and EORTC). D. As C but differential gene set enrichment. E. Heatmap showing the Jaccard similarity between the validated gene sets from D. F. The performance of different regressors built to predict TRS from primary GBM gene expression data (grey; training points, colour; blind test centres, See Supp.Table.1). G. Mutational burden in Up versus Down responder primary GBM tumours per cohort (Discovery n = 23) and filtering for different functional impacts. H. The highest ranked odds ratios of being an Up responder given mutations in specific genes across both cohorts (top 20 Fisher p with 95% CI, diamonds capped at ±3). I. The overlap between genes within odds ratios with uncorrected p<0.05 across cohorts (top) and the mutations within the only gene reaching this threshold in both cohorts: SRSF11 J. The JBS axis score was derived so that the change in score between two longitudinal GBMs equates to that patient’s TRS.

### The direction of transcriptional reprogramming is not reliably predicted at primary diagnosis

We collated all available bulk RNAseq data for paired GBMs with tumour purity >40%, forming two independent datasets: the Discovery and EORTC cohorts (Supp.Tables.1 and 2)(7,8). Responder subtype is a binary abstraction that likely discretises a continuous biological response, reducing the information available to develop a prediction algorithm. We, thus, developed a continuous variable that quantifies post-treatment changes in JBS gene expression: the transcriptional response score (TRS). TRS showed no association with continuous clinical variables (age at diagnosis, surgical interval, overall survival, MGMT expression or purity in the primary tumour) nor with categorical ones (gender, tumour location, molecular subtype and MGMT promoter methylation status) (Supp.Fig.1).

We next examined whether primary GBM molecular profiles can predict TRS. Differential expression analysis showed that very few individual genes were significantly associated with TRS in both cohorts (Fig.1C). In contrast, gene set enrichment analysis identified 145 significant pathways shared across cohorts (Fig.1D and Supp.Table.3) with low pairwise Jaccard similarity scores confirming that this was not owing to shared genes amongst the enriched sets (Fig.1E). Inspecting the validated gene sets shows that Down responder primary GBMs are enriched in mature brain, neuronal circuitry, and structural programmes, whilst Up responder primary GBMs are enriched for neurodevelopmental progenitor signatures, cell cycle activity and DNA/RNA metabolism (Supp.Table.3). Notably, these programmes correspond to those induced through treatment in the opposite responder subtype (Fig.1B). We proceeded to develop machine-learning regressors using single-sample gene-set enrichment scores and scFoundation embeddings. Despite extensive optimisation of feature selection, model architecture, and hyperparameters, performance was weak across both the training data and blind test datasets (Supp.Table.4 and Fig.1F), suggesting that responder subtype cannot be reliably predicted from primary GBM transcriptomes.

Analysis of variant calls likewise revealed no significant differences in mutational burden between responder subtypes (Fig.1G). Gene-level analyses identified no variants that robustly segregated responder subtypes, and no associations remained significant after correction for multiple testing (Fig.1H). Variants in RNA splicing factor *SRSF11* did, however, show nominal significance (p<0.05) in both cohorts, with Down responders exhibiting variants of greater predicted functional impact despite occasional variants also being observed in Up responders (Fig.1I).

Together, these findings suggest that neither responder subtype nor TRS can be predicted from clinical characteristics or the molecular profile of the primary GBM.

### Individual bulk GBM tumours can be placed on a transcriptionally defined axis

In our earlier study, we proposed that responder subtypes reflect SoC-induced movement of GBM cells along the proneural-to-mesenchymal (PN-MES) phenotypic axis described by Wang et al. (Fig.1B)(7,10,16). We therefore defined a JBS axis based on JBS gene expression and their consistent post-SoC dysregulation (see Methods), allowing individual tumours to be positioned within a common transcriptional framework. Within this framework, the difference between the JBS axis scores of paired primary and recurrent GBMs defines the TRS (Fig.1J). This enables direct comparison of primary and recurrent tumours as well as single-cell, spatial transcriptomic, and multi-region datasets within a shared reference space.

### The transcriptionally defined axis corresponds with bulk tumour phenotypes

We assessed how the JBS axis compares with the PN-MES axis defined by Wang et al. We observed significant correlation in both the Discovery (r=-0.46,p=8.4E-8) and EORTC (r=-0.42, p=2E-6) datasets, with lower JBS axis scores corresponding to mesenchymal, and higher JBS axis scores with proneural, cancer states (Fig.2A). GSEA showed that low JBS axis scores, in both primary and recurrent tumours, consistently associate with mesenchymal malignant states, stress and hypoxia responses, extracellular matrix remodelling, inflammation and immune activation, reactive astrocytes, angiogenesis, and histological features of microvascular proliferation and pseudopalisading necrosis (Supp.Table.5). High JBS axis scores associate with neuronal circuitry pathways, synaptic function, oligodendrocytes, malignant NPC and OPC states, and histological features of infiltrating tumour and the leading edge (Supp.Table.5).

**Figure 2.**
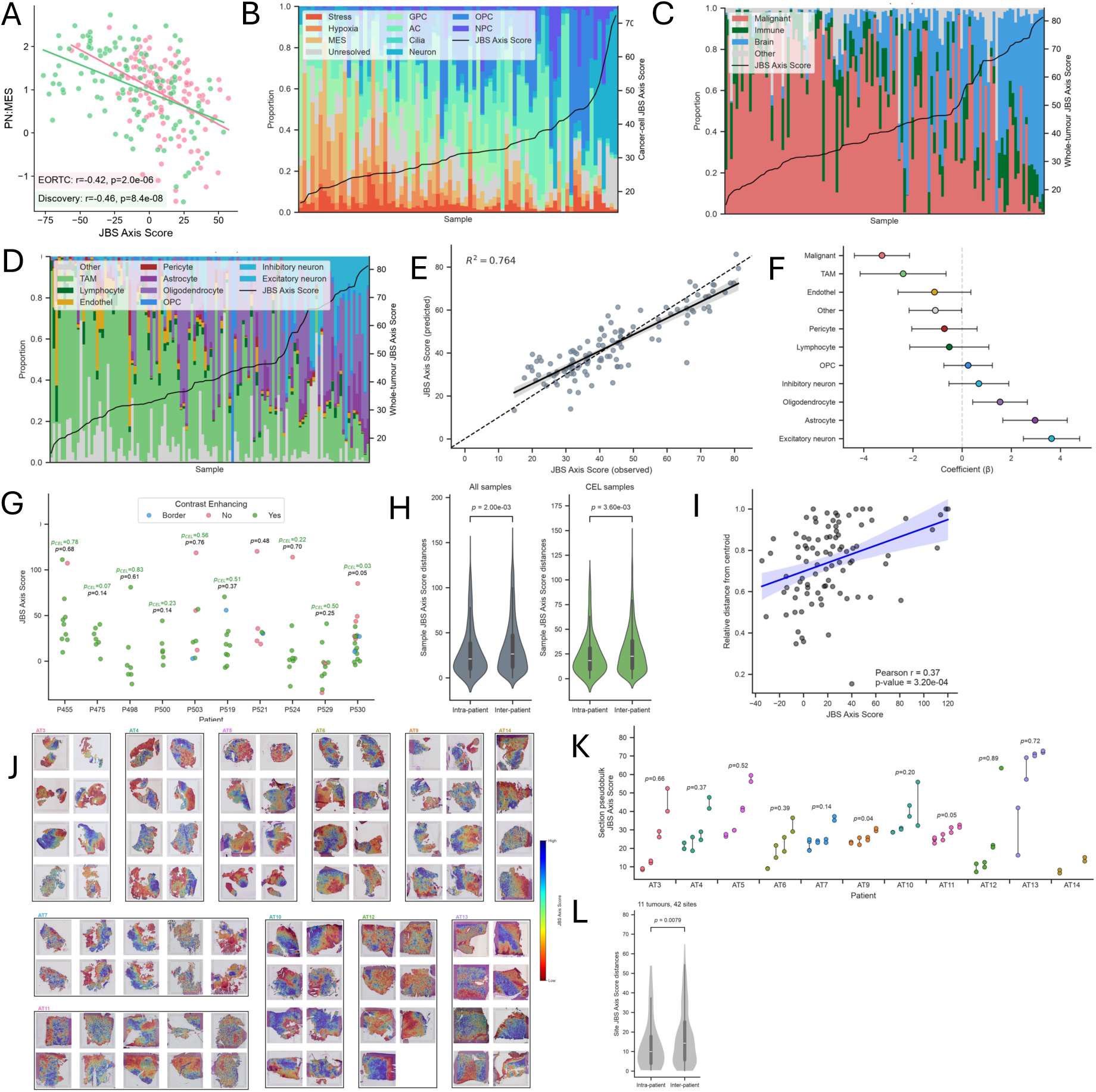
A. The correlation between the JBS axis score and the proneural to mesenchymal (PN:MES) axis score from Wang et al, 2022. **B.** The neoplastic cell type composition of GBM tumours ordered by their JBS axis score. **C.** The cell group prevalence in tumours ordered by their JBS axis score. **D.** The tumour microenvironment cell type composition of GBM tumours ordered by their JBS axis score. **E.** The performance of a linear regression model predicting JBS axis score from cell type prevalence. **F.** The coefficients each cell type received in the linear regression model in E. **G.** The JBS axis score given to different MRI-guided surgically removed regions of GBM tumours. **H.** The inter- and intra-patient distributions of JBS axis scores shown in G, for all samples (left) and only those from contrast-enhancing regions of the lesion (CEL; right) . **I.** Dotplot showing the MRI-guided regions according to their JBS axis score and distance from the tumour centroid **J.** GBM tumour sections from 12 patients that have undergone Visium spatial transcriptomics, coloured by the JBS axis score for each spot. **K.** Sections from J plotted according to their mean JBS axis score. Lines connect data from sections of the same tumour site. **L.** The inter- and intra-patient distributions of JBS axis scores shown in K.

We hypothesised that the JBS axis aligns with the PN-MES axis derived from single-cell GBM data, while extending it to bulk tumours containing non-neoplastic populations. To test this, we analysed single nuclei data and examined cellular composition across the JBS axis(9). JBS axis scores derived from cancer cell only pseudobulks increased with higher proportions of proneural cell states whereas higher mesenchymal compositions result in lower JBS axis scores (Fig.2B). JBS axis scores derived from whole tumour pseudobulks increase progressively with higher proportions of brain-resident populations (Figs.2C), with the prevalence of neurons being particularly associated with higher scores (Figs.2D). Linear regression analysis using cell-type prevalences as predictor variables showed that cellular composition accounted for 76% of the variation in JBS axis scores across whole-tumour pseudobulks (R² = 0.76, Fig.2E). Of note: within this model, healthy brain- resident cells contribute to positive axis score predictions whereas immune and vascular cells contribute towards negative scores (Fig.2F).

This establishes that the JBS axis captures reproducible biological variation with malignant and stromal cell populations occupying distinct positions along a proneural to mesenchymal related spectrum. The JBS axis score is thus expected to vary intratumourally across TME niches. To confirm this, we analysed published data from spatially distinct regions of multiple GBM tumours, acquired via MRI-guided 3D sampling(17). JBS axis scores for these multi-region samples displayed substantial intra-tumour dispersion, with individual patients showing both strongly positive and strongly negative values (Fig.2G). Samples were annotated as having being removed from the contrast-enhancing component of the lesion (CEL) or the non-enhancing component (non-CEL). Non-CEL samples often showed high JBS axis scores, consistent with infiltrated brain parenchyma, although some exhibited the lowest scores, likely reflecting poorly vascularised hypoxic or necrotic tumour regions (Fig.2G)(18). Despite substantial intratumoural heterogeneity, samples from the same tumour clustered more closely along the JBS axis than samples from different patients, both across all regions and within CEL-only subsets (Fig.2H; p=2.0E-3 and 3.6E-3 respectively). Using the sample coordinates provided with these data, we found a modest but significant correlation between JBS axis score and the relative distance of the sample from the centre of the tumour (Fig.2I; Pearson r=0.37; p=3.2E-4).

We next analysed Visium spatial transcriptomics data of primary GBM samples from De Jong et al.(19). Quantification of the JBS Axis score in each spatially resolved spot revealed clear spatial patterning in all tumour slices (Fig.2J). Using the JBS axis score derived from pseudobulking all spots per section, we could independently reassess how scores alter across tumour regions. We found clear intratumoural heterogeneity (Fig.2K). However, as per the MRI-guided samples, JBS scores were significantly more variable across patients than within single tumours (Fig.2L; p=7.9E-3).

Cumulatively, these data confirm that the transcriptionally defined JBS axis corresponds with bulk GBM tumour phenotypes consisting of specific cancer and microenvironmental cell types.

### GBM tumours phenotypically polarise post-SoC

One interpretation of the impact of TME elements shown in Fig 2, is that that GBM patients do not stratify according to their response to treatment, but rather that the bidirectional transcriptional response phenomenon is simply the effect of sampling different tumour microenvironments at primary and recurrence. To investigate this we plotted JBS axis score distributions in primary and recurrent tumours separately and found that they differed markedly. Gaussian mixture modelling shows that primary GBM scores form a single peak centred near zero, whereas recurrent tumours display a bimodal distribution polarised towards either end of the axis (Fig.3A). We therefore tested whether gene expression profiles could distinguish primary from recurrent GBMs and whether TRS could be predicted more accurately from recurrent than primary tumours. Both were true: a support vector machine classified primary and recurrent GBMs with 90% accuracy, 92% sensitivity, and 86% specificity, while a random forest trained on recurrent tumours predicted TRS in blind test data with an R² of 0.60 compared with 0.25 when trained on primary tumours (Supp.Tables.4 and 6). Together, these findings indicate that recurrent GBMs are polarised along the phenotypic axis, raising the question of whether this reflects intrinsic reprogramming, altered cellular composition, or both.

**Figure 3.**
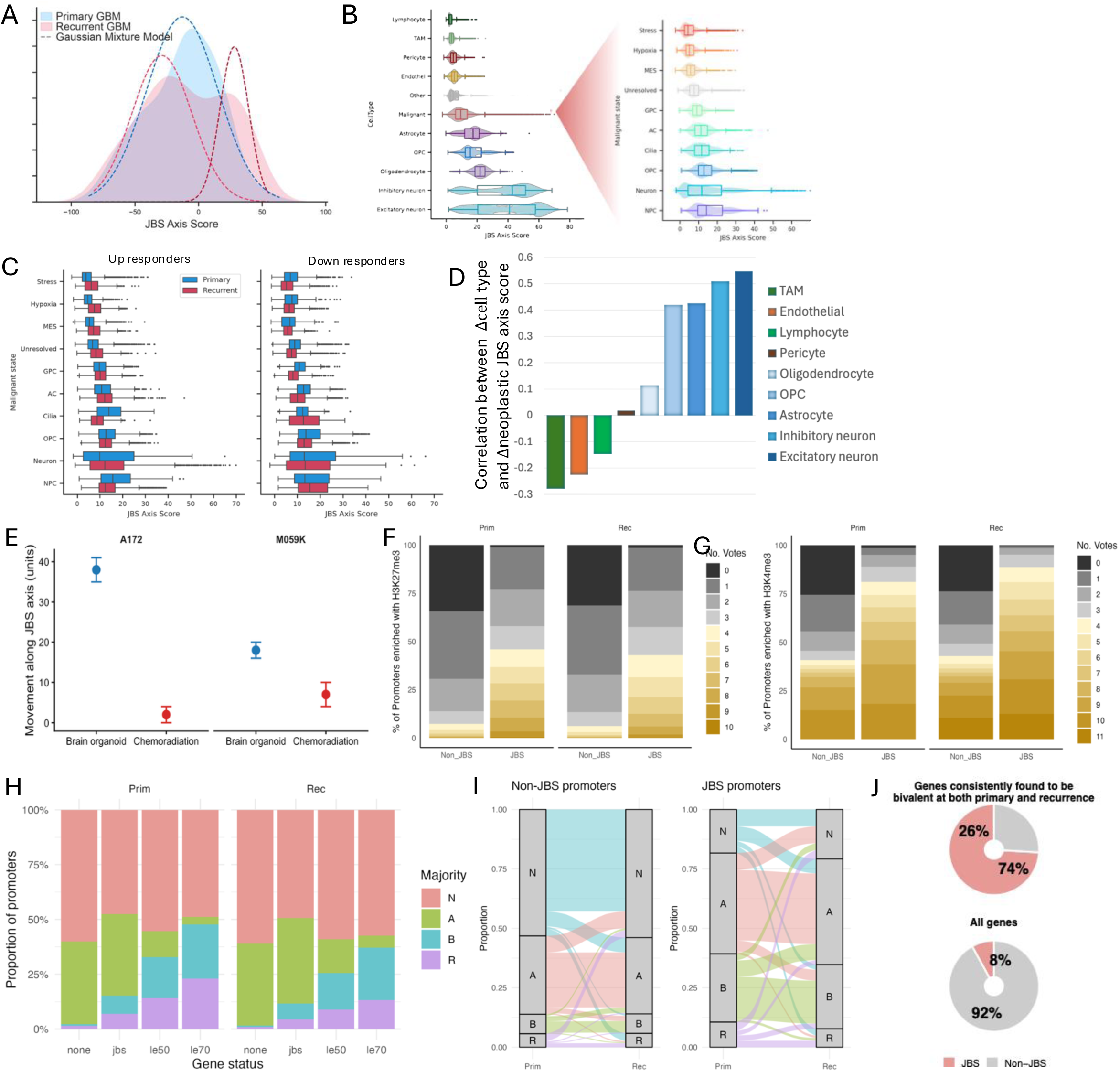
A. JBS axis score distributions for primary versus recurrent GBM tumours, overlaid with the Gaussian mixture models that best fit each distribution. **B.** JBS axis score distributions for single neoplastic and microenvironment cells (left) and for neoplastic GBM cells subtypes only (right) **C.** JBS axis score distributions for single neoplastic GBM cells in primary versus recurrent tumours, in Up and Down responders separately. **D.** Correlation between the change in TME cell type from primary to recurrence and the change in JBS axis score of the neoplastic cells in the same tumour **E.** The magnitude of change in JBS axis score in two GBM cancer cell lines (A172 and M059K) when added to a brain organoid versus being subjected to a single dose of chemoradiation. **F.** The proportion of promoters (separated into JBS gene promoters and Non-JBS) classed as being enriched for H3K27me3 in 0-10 primary or recurrent patient GBM tumours **G.** As per F but for 0-11 primary and recurrent tumours successfully profiled for H3K4me3. **H.** The proportion of promoters classed, by a majority across patient samples, as having neither histone mark (N) the activating H3K4me3 mark alone (A) the repressive H3K27me3 mark alone (R) or having both marks and being, thus, bivalent (B) in primary and recurrent GBM tumours. Promoters are separated into those from Non-JBS genes and JBS genes, the latter of which are further subsetted into those in the leading edge of at least 50% of the patients profiled (LE50) or 70% (LE70) as per Tanner et al, 2024. **I.** Sankey plots showing the proportion of histone mark state changes between paired primary and recurrent tumours in both Non-JBS and JBS gene promoters **J.** (Top) Promoters classed, by majority, as bivalent in both the primary and recurrent tumour broken down by JBS status (bottom) all promoters similarly broken down by JBS status.

### Phenotypic polarisation corresponds to changes in tumour microenvironment

To assess how post-SoC changes in cell abundance influence polarisation, we revisited the longitudinal GBM single nuclei data to investigate the arrangement of different GBM tumour cell types on the JBS axis. Single nuclei and cell data present specific technical challenges for JBS axis scoring and require additional processing steps. Consequently, whilst single-cell axis score distributions retain interpretability in terms of relative ordering and magnitudes, they are dataset-specific and almost always >0. Fig.3B shows the distribution of GBM tumour cell types along the JBS axis. Non-malignant cells are ordered, from low to high scores, from immune and endothelial to normal brain, with neurons scoring most highly. Of note, when separating the malignant fraction into the GBM cell types defined in Nomura et al(9). we see a clear gradient, with mean scores transitioning from mesenchymal-like states to proneural-like states in an order that mirrors the malignant state transitions reported in that study. Further focusing on the malignant states but separating primary from recurrence we see shifts in axis score distributions within the window of that cell’s annotation. Interestingly, these shifts in score distributions from primary to recurrence within each malignant cell type are most often in opposing directions for Up and Down responders (Fig.3C). Whilst it is not possible to ascertain if, and how often, neoplastic cell movement along the JBS axis results in a new cell type being assigned, these opposing shifts in scores within cell type do suggest phenotypic adaptation of the neoplastic cells themselves. Furthermore, changes in non-malignant cell abundance between paired primary and recurrent GBMs correlated with neoplastic-cell JBS axis polarisation (Fig.3D). Increased lymphocyte abundance was associated with negative neoplastic polarisation (r = −0.27, p = 0.05), whereas increased abundance of OPCs (r = 0.42, p = 0.002), astrocytes (r = 0.43, p = 0.002), inhibitory neurons (r = 0.51, p = 1.6 E-4) and excitatory neurons (r = 0.55, p = 4.0 E-5) was associated with positive neoplastic cell polarisation.

These findings indicate that recurrence-associated GBM polarisation reflects enrichment of either a normal brain/leading-edge microenvironment with neoplastic cells polarised to more proneural phenotypes (Up responders) or an immune-rich, tumour-dense microenvironment with neoplastic cells polarised to more mesenchymal phenotypes (Down responders). This may result from selective expansion of unresected cells already primed to a specific niche, or from adaptive reprogramming of residual tumour cells to the post-surgical microenvironment, potentially mediated by JBS gene dysregulation. To investigate this, we proceeded to inspect whether established, phenotypically homogeneous GBM cell lines move along the JBS axis following exposure to chemoradiation and to a new TME.

### GBM cancer cells change expression of JBS genes in response to treatment and changes in TME

In our initial study, 3D spheroids derived from established (A172, M059K) and patient-derived GBM cell lines altered JBS gene expression in response to temozolomide and irradiation despite the absence of microenvironmental cells, although the effect sizes were considerably smaller than those observed in patients(7). To extend these findings, we allowed A172 and M059K spheroids to invade brain organoids and establish a brain-like tumour microenvironment (TME). Following RNA sequencing of isolated neoplastic cells, we found that the addition of a brain-like TME altered JBS axis scores substantially more than chemoradiation. In M059K cells, brain organoid invasion shifted the JBS axis score by 18 ± 2 units compared with 7 ± 3 units following chemoradiation, while in A172 cells the corresponding shifts were 38 ± 3 and 2 ± 2 units, respectively (Fig.3E). Given the homogeneity of these well-established cell line models, these findings suggest that GBM cells adapt transcriptionally to both chemoradiation and, more profoundly, to changes in their microenvironment, as may occur following surgery. As JBS dysregulation appears central to this adaptive response, it may represent a therapeutic vulnerability warranting further investigation. We, thus, sought to understand how these genes are epigenetically regulated in primary and recurrent GBM.

### JBS gene promoters are significantly more likely to be bivalent, and retain bivalency through treatment

We previously showed that JBS gene promoters are enriched for bivalent chromatin, marked by both H3K27me3 and H3K4me3, in primary GBM(20). To determine whether treatment-associated phenotypic adaptation is accompanied by resolution of this state, we profiled both histone marks in 13 fresh frozen paired GBM tumours. After QC, 10 primary and 11 recurrent GBMs had H3K4me3 profiles (Supp.Table.7), giving 9 matched pairs, and 10 primary and 10 recurrent GBMs had H3K27me3 profiles, yielding 8 matched pairs (Supp.Table.8); with 8 pairs successfully profiled for both marks. As expected, JBS promoters in primary tumours were significantly enriched for H3K4me3 (OR=6.22, Fisher’s P=0), H3K27me3 (OR=10.80, Fisher’s P=0) and bivalency (OR=22.27, Fisher’s P=0) relative to non-JBS genes (Fig.3F-G; left). Strikingly, equivalent enrichment was observed in recurrent tumours for H3K4me3 (OR=10.38, Fisher’s P=0)), H3K27me3 (OR=11.42, Fisher’s P=0) and bivalency (OR=23.50; Fisher’s P=0) (Fig.3F-G; right). Enrichment increased further when analysis was restricted to the JBS genes most consistently dysregulated through treatment (Fig.3H). This suggests that promoter bivalency is not resolved post-treatment. Indeed, among paired samples, 18% of promoters classified as bivalent at either timepoint retained bivalency at both, with persistence fourfold more likely at JBS than non-JBS promoters (Fig.3I). Of the 772 promoters classified as bivalent in both primary and recurrent tumours, 573 (74%) were JBS gene promoters(Fig.3J; top), despite JBS genes comprising only 8% of all genes (Fig.3J; bottom). Functional analysis of the 772 consistently bivalent promoters revealed strong enrichment for neuronal and synaptic pathways, developmental homeobox transcription factors, and PRC2-associated gene sets, including genes marked by bivalent chromatin in stem cells. Together, these findings suggest that phenotypic plasticity enables adaptation to treatment in GBM, facilitated by Polycomb-mediated retention of bivalency within a core set of neurodevelopmental genes.

## Discussion

We previously showed that patients with GBM stratify according to the bidirectional transcriptional reprogramming of their tumour in response to standard of care (SoC) treatment (surgical resection and subsequent chemoradiation with Temozolomide)(7). Herein we showed that, whilst this finding extends to additional independent datasets (Fig.1A), it is not possible to predict responder subtype from a patient’s primary tumour (Figs.1F-H and Supp.Table.4). This agrees with findings from a recent single nuclei profiling study that concluded that longitudinal GBM trajectories cannot be predicted(2).

We proceeded to investigate whether the genes involved in the consistent transcriptional response to SoC (JBS genes, so called because they contain JARID2 binding sites in their promoter) could be used to form a framework by which individual tumours, and their subsequent bidirectional programming, could be better understood. This led to the development of the JBS axis. Individual GBM tumours, and even single cells, can be positioned on this axis based on their expression of JBS genes and the consistent post-SoC reprogramming observed therein. A patient’s responder subtype is then defined by the movement between the primary and recurrent tumour along the JBS axis. Using bulk and single cell longitudinal datasets we were able to show that a tumour’s position on the JBS axis aligns with GBM ecosystems and the cell types that define them (Figs.2A-D). Such definitions have been honed over 15 years of molecular profiling of GBM tumours, leading to distinct but complementary nomenclatures(9,16,21–24). Figure 4 illustrates how these nomenclatures align with each other and the JBS axis.

**Figure 4.**
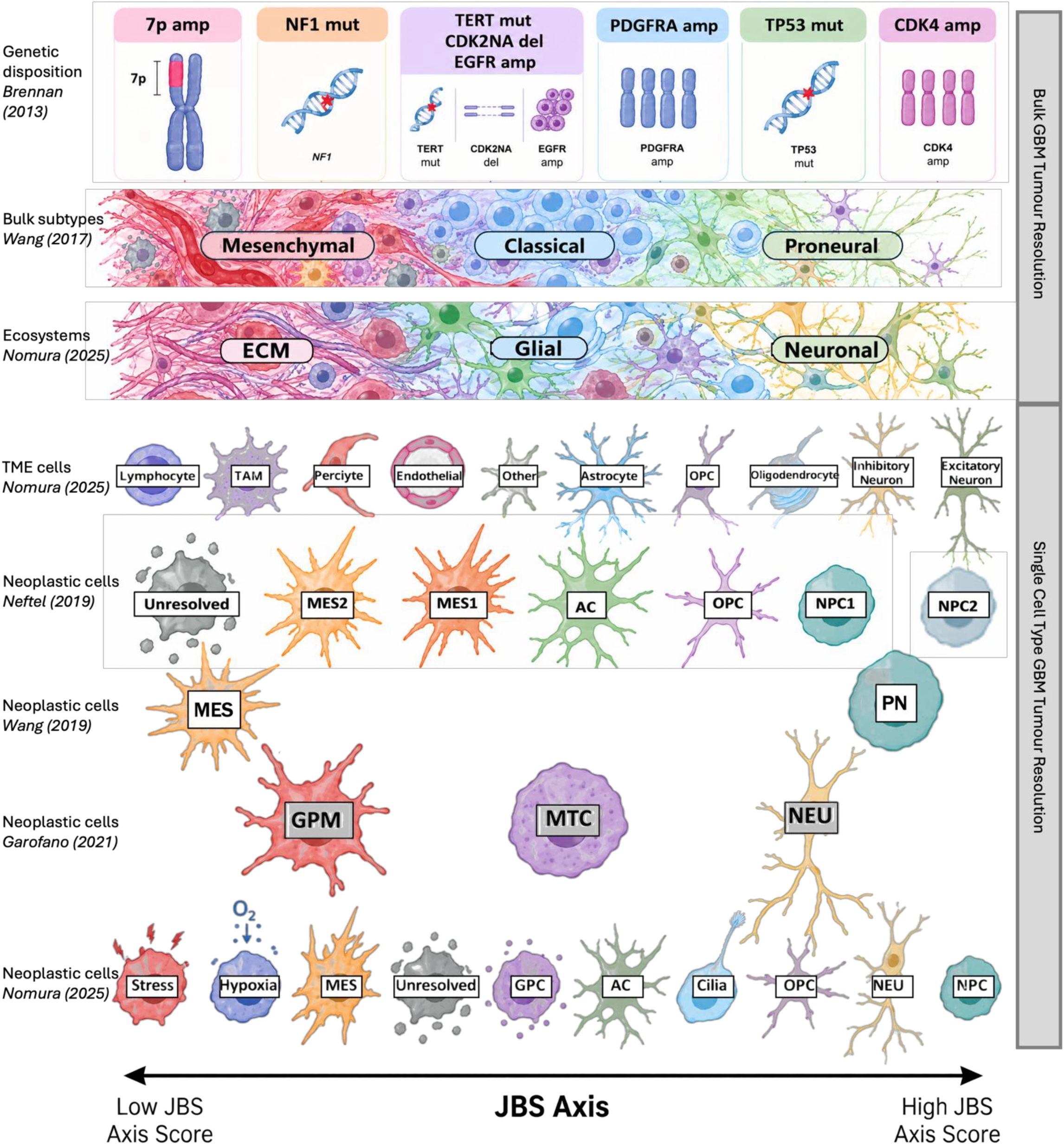
An illustration to place the JBS axis in the context of consolidated findings on bulk, and single cell, subtypes within glioblastoma brain tumours over the last 15 years. This framework highlights new avenues to understand, and therapeutically target, the ability of neoplastic GBM cells to phenotypically adapt to standard of care treatment, and the change in TME that it causes

Importantly, our findings show that recurrent tumours are polarised towards either end of the JBS axis (Fig.3A), consistent with a previous study indicating that neoplastic cells in recurrent GBM polarise towards either the proneural or mesenchymal phenotype(10,25). Recurrent GBMs have been shown to contain fewer neoplastic cells and much greater interaction with the surrounding non-malignant TME cells (2,26). Our results show that the prevalence and types of cells in the TME directly impact the JBS axis score (Figs.2E-F), and changes within the TME post-SoC influence the change in axis score within neoplastic cell populations (Fig.3D). This suggests that JBS axis polarisation is, thus, caused by the presence or influx of TME cells in the post-surgical cavity, with the dominant cell type (immune versus normal brain) determining the direction of polarisation. Using in vitro experiments, we further showed that changing the TME of neoplastic cells alters their JBS axis score significantly more than chemoradiation alone, suggesting that the cancer cells themselves plastically adapt (Fig.3E). This explains why the responder subtype and longitudinal trajectories cannot be predicted for a primary GBM: they are caused by cellular adaptation to unforeseeable alterations in the TME in response to treatment as well as to effect of chemoradiation on the cancer cells themselves.

The JBS axis score is not a conventional transcriptional state score constructed from genes that define a particular phenotype. Rather, JBS genes were selected because they are consistently dysregulated following standard-of-care treatment. Consequently, the score captures not only the tumour’s current transcriptional state, but also how closely that state aligns with the dominant recurrence-associated reprogramming trajectory. Biologically, the axis therefore represents more than a phenotypic spectrum; it positions tumours within a transcriptional landscape shaped by adaptation to treatment-induced selection pressures. This representation is validated by the faithfulness with which the JBS axis aligns to defined GBM ecosystems (Fig.4), and how changes in the score capture the ecosystem remodelling that is now known to characterise GBM evolution through SoC. Importantly, this suggests that the capacity to move between transcriptional states, rather than occupancy of a particular state, may be central to treatment adaptation and therefore represent a more broadly applicable therapeutic vulnerability.

Our results, thus, show that the genes defining the JBS axis are involved in adaptive reprogramming of GBM tumours, and that control of their dysregulation may provide an effective therapeutic opportunity. We, therefore, sought to further characterise how JBS genes are epigenetically regulated. We found that JBS gene promoters are significantly enriched in bivalent chromatin, harbouring both the active H3K4me3 and the repressive H3K27me histone marks, in both primary and recurrent tumours (Figs.3F-H). Furthermore, we found that bivalency is retained through treatment, suggesting that its resolution is not the mechanism driving bidirectional dysregulation of JBS genes (Figs.3I-J). Of note, genes that retain bivalency post-SoC are enriched for processes of (neuro)developmental cell-state regulation.

Bivalent chromatin was originally proposed to maintain developmental genes in a repressed but transcriptionally poised state in embryonic stem cells(27). Subsequent studies have challenged the view that bivalency universally predicts rapid gene activation, suggesting instead that bivalent domains contribute to developmental competence and epigenetic flexibility(28,29). Crucially, JARID2 is required for the transcriptional priming of bivalent genes in pluripotent stem cells, a process that is essential for maintaining their capacity for multi-lineage differentiation(15).

Our results suggest that persistent bivalency, mediated through JARID2-associated Polycomb activity at JBS genes, provides a plausible molecular mechanism through which GBM cells maintain the ability to adapt to, and transition between, treatment-induced ecosystem states. Rather than resolving developmental programmes during tumour evolution, recurrent GBMs appear to retain a substantial reservoir of poised neurodevelopmental loci, enabling continued plasticity and adaptive power. In this way, understanding and targeting the mechanisms by which bivalency is established and retained at these loci may present a therapeutic opportunity. This work supports two landmark single cell multiomics papers published recently, which also suggest that limiting the capacity for transcriptional adaptation, irrespective of its direction, could provide a therapeutic strategy to constrain recurrence(19,30).

We do note that, as histone marks were profiled separately in bulk tumour tissue, their co-occurrence at individual promoters within the same cells cannot be confirmed. Some apparent bivalency may therefore reflect distinct chromatin states across heterogeneous cell populations, warranting validation using cell-resolved approaches.

In addition, future work should identify the epigenetic regulators and transcription factors responsible for maintaining this plasticity and executing JBS gene reprogramming, including whether these differ between responder subtypes. Although JARID2 is an obvious candidate, its functional contribution to GBM adaptation remains to be established. Identifying and targeting these mechanisms alongside SoC treatment may offer a means to prevent or delay the adaptive responses that enable tumour recurrence.

## Methods

All statistical analyses were performed in R and Python. Unless otherwise stated, tests were two-sided, and multiple-testing correction was applied using the Benjamini-Hochberg procedure. Visualisations were generated using ggplot2, seaborn and matplotlib.

### RNA sequencing datasets and preprocessing

Bulk RNA-seq, single-cell RNA-seq (scRNA-seq), and spatial transcriptomic datasets were compiled from published studies and newly generated samples (Supplementary Tables S1-S2). These included longitudinal paired primary and recurrent IDH-wildtype glioblastoma (GBM) cohorts together with additional de novo IDH-wildtype GBMs used for dimensionality reduction analyses. Forty-two additional longitudinal formalin-fixed paraffin-embedded (FFPE) GBM samples were processed and sequenced as previously described. Where available, raw sequencing data were reprocessed using a standardized pipeline; otherwise, published count matrices were obtained. For scRNA-seq data, quality control was performed using Scanpy. Cells expressing ≥200 genes and <3% mitochondrial transcripts were retained, and doublets were removed computationally. Cell-type annotation was adopted from the original studies. For spatial datasets, quality-controlled count matrices were obtained from study authors after removal of low-quality spots and regions outside tissue boundaries. Samples were restricted to patients with de novo IDH-wildtype GBM treated with standard-of-care therapy and experiencing local recurrence. Tumours with estimated purity <40% were excluded using GBMPurity. Patients were divided into independent Discovery (n=62) and EORTC (n=58) cohorts that were analysed separately for validation purposes. For machine-learning analyses, DEA and GSEA (for Figure 1), 32 Discovery cohort patients processed at a single institution were combined with the EORTC cohort to create a training set (n=90), while the remaining Discovery cohort patients constituted an independent test set (n=30).

### Transcriptomic analyses

Cellular deconvolution was performed using GBMdeconvoluteR(31) and molecular subtype assignment using the three-class SubtypeME classifier. Differential expression analyses were conducted with DESeq2. For analyses associated with transcriptional response scores (TRS) or JBS-axis scores, differential-expression statistics were ranked by sign(log2 fold-change) × −log10(adjusted P value). Gene-set enrichment analysis (GSEA) was performed using preranked gene lists and curated pathway collections from KEGG, Reactome, WikiPathways, and a previously described GBM-specific pathway resource. Gene sets containing 15-5,000 expressed genes were retained.

### RNA-seq variant calling

RNA-seq somatic variant calling was performed using a Nextflow-based workflow. Reads were aligned to GRCh38 using STAR and processed according to GATK best-practice recommendations, including duplicate removal, splice-aware preprocessing, and base-quality recalibration. Somatic variants were identified using Mutect2 with population germline resources and genomic blacklist filtering. To minimise false-positive calls arising from RNA-seq artefacts, variants overlapping known germline polymorphisms, RNA-editing sites, and pseudogene regions were removed. Remaining variants were annotated using Ensembl Variant Effect Predictor. Gene-level enrichment analyses were performed using odds ratios calculated with maftools.

### Development of the transcriptional response score

Patients were assigned responder subtypes as previously described based on the enrichment direction of the JBS gene signature between matched primary and recurrent tumours. The transcriptional response score (TRS) was defined as the first principal component derived from principal-component analysis of JBS-gene log2 fold-change values in the Discovery cohort.

### Development of predictive classifiers

Only protein-coding genes present across all datasets were retained. To identify transcriptomic representations associated with TRS, four dimensionality-reduction approaches were evaluated: non-negative matrix factorisation (NMF), contractive autoencoders, scFoundation embeddings, and single-sample gene set enrichment analysis (ssGSEA). Feature performance was assessed according to association with TRS and cross-validated predictive accuracy. scFoundation- and ssGSEA-derived representations demonstrated the strongest performance and were subsequently advanced for model development. ElasticNet and Random Forest regression models were trained using Bayesian hyperparameter optimisation and repeated cross-validation within the training cohort. Final models were retrained on the complete training dataset and evaluated in the held-out test cohort. Predictive performance was assessed using coefficient of determination (R²) and root mean squared error.

### Calculation of JBS and PN-MES axis scores

To enable projection of independent datasets, JBS-axis scores were calculated by multiplying log2-transformed JBS-gene expression values by the principal-component loading vector derived from the Discovery cohort and summing across genes. These fixed loadings were applied to all bulk, single-cell and spatial datasets to ensure comparability between analyses.

For datasets available only as TPM-normalised data, scores were linearly transformed using an empirically derived offset to align with FPKM-derived values. PN-MES axis scores were calculated similarly using published principal-component loadings.

### Gaussian mixture modelling and polarisation analyses

To quantify the presence of discrete cellular states, Gaussian mixture models were fit to JBS-axis score distributions. Model performance was assessed using the Bayesian Information Criterion (BIC), and evidence for bimodality was determined by comparing one- and two-component models. Additional measures of population segregation included Ashman’s D, posterior assignment entropy, and silhouette coefficient. Confidence intervals were estimated using bootstrap resampling.

### Compositional and state-transition analyses

Cell-type abundances were expressed as proportions of total cells and transformed using centred log-ratio (CLR) transformation before statistical modelling. Associations between composition and transcriptomic state were assessed using ordinary least squares regression. To examine treatment-associated state dynamics, JBS-axis scores were calculated for individual cells. Median values were determined separately for primary and recurrent tumours within each patient and cell population. Changes in median JBS-axis score across treatment were correlated with corresponding alterations in CLR-transformed cellular composition.

### Glioma-brain organoid model

Human ReNcell-VM neural stem cells were cultured under non-adherent conditions to generate neural spheroids and differentiated for 7 days. GFP-labelled A172 and M059K glioblastoma spheroids were generated separately and combined with neural spheroids to create glioma-brain organoids (GLIBOs). Following co-culture, organoids were dissociated, GFP-positive tumour cells were isolated by fluorescence-activated cell sorting, and RNA was extracted for sequencing. Libraries were prepared using standard Illumina-compatible protocols and sequenced to approximately 50 million paired-end reads per sample.

### Chromatin profiling and epigenomic analysis

Fresh-frozen longitudinal GBM tissues were processed for CUT&RUN profiling of H3K4me3 and H3K27me3 by The Epigenomics Profiling Core at MD Anderson Cancer center as previously described but with some modifications(32). Nuclei were isolated, immobilised on Concanavalin A-coated beads, and incubated with IgG (Millipore), H3K4me3 (EpiCypher) and H3K27me3 (Diagenode) antibodies overnight at 4°C. Next day, targeted chromatin digestion was achieved by pAG-MNase binding and incubation with CaCl_2_ at 4°C, followed by CUT&RUN DNA purification. Libraries were prepared using standard Illumina workflows and sequenced on a NextSeq 2000 instrument. Matched H3K4me3 ChIP-seq datasets were additionally generated for selected samples. Sequencing reads were quality controlled, adapter trimmed, aligned to GRCh38, and filtered to retain uniquely mapped read pairs. Histone-mark enrichment at promoters was assessed relative to matched IgG controls. Promoters were defined as ±1 kb around annotated transcription start sites. Samples failing quality-control metrics based on promoter enrichment and concordance with published GBM chromatin profiles were excluded.

## Author Contributions

LFS conceived the study with input from MT, and secured funding with input from JP. MDJ, MDC and ARB coordinated patient recruitment, tissue acquisition, and clinical data collection. MT, BW, and LFS performed data processing, bioinformatic analyses, and statistical analyses with support from YA, MA and UGK. LFS and AB supervised data analyses. AKJ, MF, SP and IC carried out tissue preparation and laboratory experiments. LFS, AJ, JP, TAH, MW and LM supervised lab work. LFS and MT interpreted the findings. MT prepared all figures, with input from LFS. LFS and MT drafted the manuscript. All authors reviewed and approved the final manuscript.

## Acknowledgments

This work was supported by grants from UK Research and Innovation [MR/T020504/1 and UKRI1069 to LFS, CDT grant number EP/S024336/1], The Brain Tumour Charity [ET_2023_2 10759 to LFS and JP], Yorkshire’s Brain Tumour Charity and OSCARs Paediatric Brain Tumour Charity [Joint Infrastructure funding to LFS] and Leeds Hospital Charity [A2002546 to LFS]. YA was supported by a doctoral scholarship from King Abdulaziz Medical City, Ministry of National Guard Health Affairs (MNGHA), Riyadh, Saudi Arabia. MA was supported by a doctoral scholarship from The Oman Medical Specialty Board. AKJ was partially supported by MD Anderson Cancer Center institutional support to Epigenomics Profiling Core. Tissue used in this study was accessed from: the Leeds Neuropathology Research Tissue Bank, funded by Yorkshire’s Brain Tumour Charity, OSCARs Paediatric Brain Tumour Charity and Leeds Hospitals Charity; the Sidney Driscol Neuroscience Foundation BTNW tissue bank; and from Cambridge University Hospitals NHS Foundation Trust and Imperial College Healthcare NHS Trust as part of the UK Brain Archive Information Network (BRAIN UK) which is funded by the Brain Tumour Research. We wish to thank all of the patients who donated their tissue for the use in research; we sincerely hope this work can accelerate us towards more effective treatments.

## Data Availability Statement

The data generated in this study will publicly available in the European Genome-phenome Archive (EGA) upon publication. The data analysed in this study were obtained from various repositories, listed, alongside accession codes, in Supp.Table.1.

## Code Availability Statement

The custom code used to perform the analyses and generate the figures presented in this study is publicly available on GitHub (GliomaGenomics/TRS-GBM) and through a reproducible computational capsule hosted on Code Ocean (Capsule 7000550).

## Ethics Statement

All samples were from patients that provided informed, written consent. The use of consented samples was following favourable opinion of a project-specific application by NHS Health Research Authority: South Central - Oxford A Research Ethics Committee (reference 13/SC/0509).

## Notes

### Competing Interest Statement

The authors have declared no competing interest.

