## Supplemental Figures for "Glioblastoma exhibits bidirectional adaptation to treatment through JARID2-associated developmental plasticity"

### Slide 1
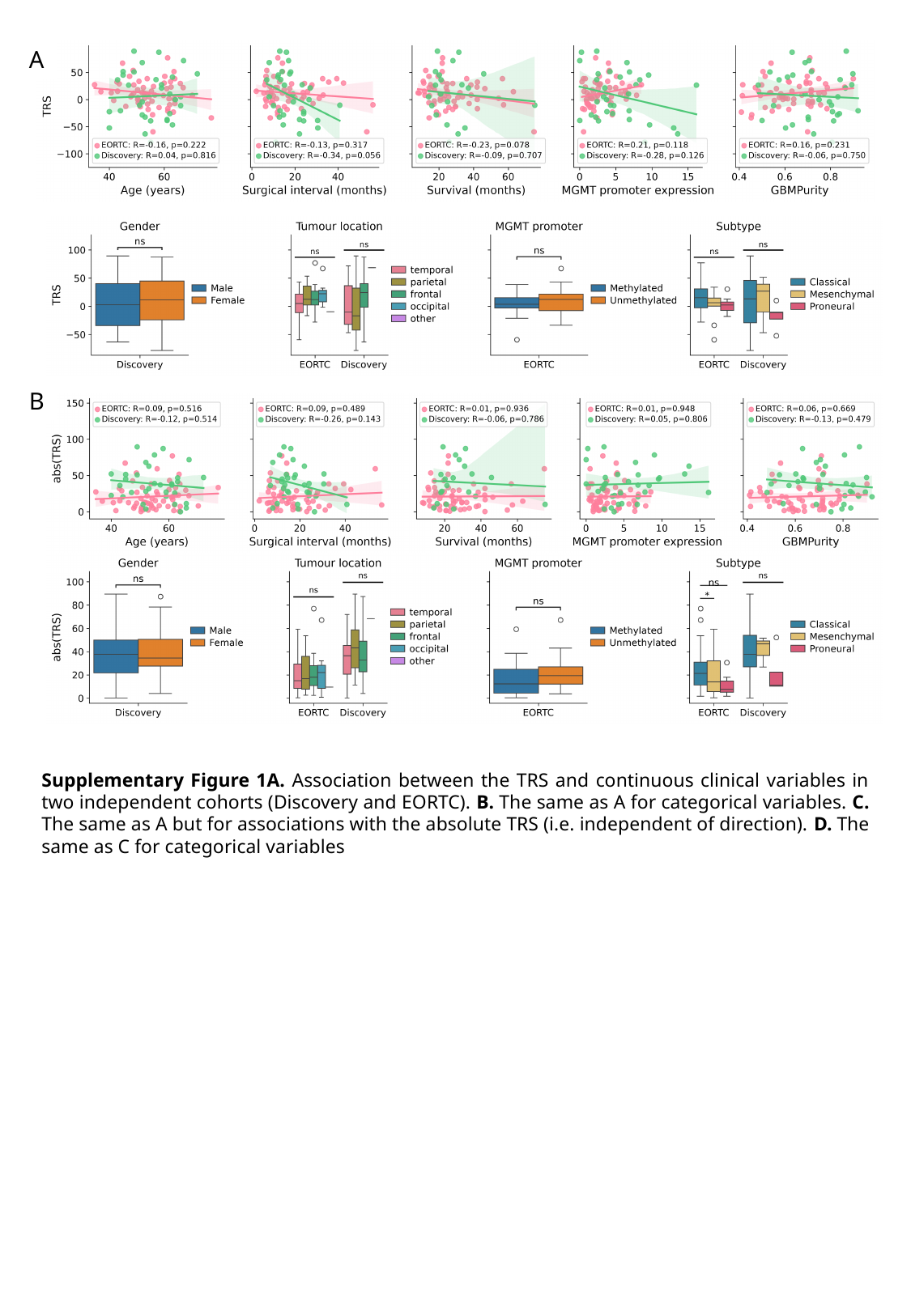

A
B
Supplementary Figure 1A. Association between the TRS and continuous clinical variables in two independent cohorts (Discovery and EORTC). B. The same as A for categorical variables. C. The same as A but for associations with the absolute TRS (i.e. independent of direction). D. The same as C for categorical variables
