## Supplemental Methods for "Glioblastoma exhibits bidirectional adaptation to treatment through JARID2-associated developmental plasticity"

**RNAseq cohorts and dataset processing**

Multiple bulk and scRNA-seq datasets were used in this study, including unmatched de novo IDHwt GBM data for dimensionality reduction purposes. All datasets are detailed with source citations and key characteristics in SuppMethodsTables1 and 2. Bulk raw sequencing or alignment data, where available, was processed through a previously described pipeline(1). This included raw RNA sequencing data created specifically for this project from 42 additional longitudinal GBM FFPE samples, processed and sequenced as previously described(1). Where raw data was not accessible, count matrices were acquired. For single-cell data, quality control procedures were implemented in scanpy (v1.11.1). Cells expressing ≥200 genes and ≤3% mitochondrial gene counts were retained. Doublets were identified and removed using scanpy’s scrublet function. Pseudobulk profiles were generated as required prior to normalisation. Cell type annotation was taken from the originating papers. For spatial data, quality-controlled, preprocessed raw count matrices for each sample, with low quality spots and those reside outside tissue boundaries removed, were obtained from the authors.

All data was normalised to either CPM, TPM and FPKM with each format used as needed for downstream analysis (see below). Longitudinal datasets were filtered for patients diagnosed with de novo IDHwt GBM who under standard treatment and had a local recurrence. All data were manipulated using Anndata (v0.10.9), scanpy (v1.11.1), and Seurat (v5.1.0). Batch correction was performed using the pyComBat-seq implementation of the ComBat-seq algorithm in the inmoose package (v0.7.3) for single sample analyses, and centred log-ratio transformation for compositional data analysis. Data was filtered for minimum tumour purity of 40% in all matched pairs according to GBMPurity(2) and separated into two cohorts: Discovery (n=62) and EORTC (n=58). These two cohorts were analysed separately for all of the downstream analysis, to enable cross-cohort validation, except for machine learning. For the latter, 32 Discovery cohort patients, all processed at the University of Leeds, were added to the 58 EORTC patients to create a training dataset of n=90. The remaining 30 Discovery cohort patients were retained as a test data set.

**RNAseq analyses**

All cellular deconvolution was performed in GBMDeconvoluteR(3) using FPKM inputs with the default GBM marker genes. Molecular subtyping was done using the 3-way SubtypeME tool in GlioVis (<http://gliovis.bioinfo.cnio.es/>) using TPM inputs. Differential gene expression analysis (DEA) was performed using DESeq2 (v1.42.1). For responder subtype prediction, genes were stratified as protein-coding and non-coding genes and the TRS target was rescaled: *TRS_scaled_=TRS/max|TRS|* and DEA performed using the design ~TRS (or ~JBS axis score where appropriate). The results were then ranked using *sign (log2 FC) ×−log_10_ padj* for input to GSEA. For GSEA, a pre-ranked list of genes was input into the fgsea R package (v1.26.0) and the fgseaMultilevel function was used with epsilon 0 andGSEA parameter of 1. Gene sets were limited to 15-5000 genes expressed above the lower quartile of non-zero expression in ≥25% of either primary or recurrent samples. 100,000 permutations were run. Gene sets were from with KEGG, WikiPathways, and Reactome, supplemented by a GBM-specific collection curated by Tanner et al.(1).

**RNAseq variant calling**

The RNA-seq variant calling pipeline was implemented and executed using Nextflow(4). Raw sequencing reads were mapped to the human reference genome (GRCh38.p10) using STAR (5). BAM files were first processed to remove PCR and optical duplicates using Picard MarkDuplicates. Following the GATK best practices workflow for raw sequencing data preprocessing, reads spanning splice junctions were processed using GATK SplitNCigarReads to split spliced alignments and assign appropriate mapping qualities(6). Subsequently, Base Quality Score Recalibration (BQSR) was performed utilizing dbSNP and the Mills and 1000G gold standard indels as known polymorphic sites to correct technical sequencing artifacts(7,8).

To mitigate library preparation artifacts and technical noise inherent to RNA sequencing data, reads were query-name sorted using Samtools and processed with fgbio ClipBam with overlapping read clipping enabled to soft-clip overlapping read segments, thereby preventing the artificial double-counting of variants within overlapping regions(9). Somatic variant calling was conducted using Mutect2, incorporating GnomAD as a population germline resource and the ENCODE blacklist to exclude regions prone to assembly errors, collapsed repeats, or structural variations that frequently generate false-positive single-nucleotide variants (SNVs) and indels due to alignment misalignment(10). To eliminate known germline polymorphisms, technical artifacts, and RNA-specific confounding events from the raw variant calls, a multi-step stringency filtering cascade was applied using Bedtools, removing known germline variants present in dbSNP (v153), variants overlapping known A-to-I RNA editing sites sourced from the Atlas database of RNA editing events in human, and variants located within GENCODE pseudogenes (11-13). Finally, functional annotation and effect prediction of the filtered somatic variants were performed using Ensembl Variant Effect Predictor (VEP) (14). MAFtools package(v2.18.0) was used to calculate odds ratios with genes ranked by *sign (log_2_ odds-ratio) ×−log_10_ nominal p-value* for input into GSEA.

**Developing the transcriptional response score (TRS)**

Binary responder subtype labels were assigned to patients as per Tanner. et al(1). Briefly, assignment is based on whether the JBS-gene set showed positive (Up responder) or negative (Down responder) enrichment within all expressed genes for that patient when ranked by the logged fold change between their primary and recurrent sample. The TRS equates to the first principal component (PC1) resulting from applying prcomp (from base R stats v4.3.0) to the JBS gene log2 fold-change gene matrix for the Discovery cohort (unless otherwise stated).

**Developing prediction classifiers**

Only protein coding genes present in all datasets were retained. All labelled and unlabelled bulk RNA-seq samples, including primary and recurrent tumours, were included in dimensionality reduction analyses, for which we tested 4 approaches: Non-negative matrix factorisation (NMF), Contractive autoencoder (CAE), scFoundation and single sample Gene Set Enrichment Analysis (ssGSEA). NMF was performed using scikit-learn v1.6.1 across ranks k=2–50k = 2–50, with ten random initialisations per rank. Model performance was assessed using Frobenius reconstruction error on training and validation datasets and compared against value-shuffled data. Data were partitioned into 80% training and 20% validation sets. Clustering stability was evaluated by consensus clustering and the cophenetic correlation coefficient. The optimal rank was selected from the cophenetic coefficient and reconstruction-error elbow plot. CAE models were implemented in PyTorch v2.6.0. Gene expression values were transformed using log_2_(x+1)\log_2_(x+1) and Z-score normalised using statistics derived from the training set. The encoder consisted of Dropout, Linear, BatchNorm and ReLU layers arranged in an input>h1>h2>latent architecture, with a symmetrical decoder terminating in a sigmoid output layer. Models minimised reconstruction MSE with an additional contractive penalty applied to the encoder Jacobian. Hyperparameters including latent dimension (8-128), hidden layer size (256-2048), dropout (0-0.5), contractive penalty (10^-10^-10^-2^), learning rate (10^-6^-10^-3^), weight decay (10^-10^-10^-4^) and batch size (8,16,32,64,128) were optimised using Bayesian optimisation in Optuna v4.3.0 over 500 trials. Training used Adam, weight decay and early stopping. Models were selected using a composite validation objective combining reconstruction loss and prediction of GBMPurity estimates using CCC((1-CCC)+MSE). scFoundation embeddings were generated using get_embedding.py. Bulk RNA-seq data were restricted to compatible genes and processed in half precision to reduce memory requirements. Max pooling was applied to obtain a fixed embedding vector per sample. Pathway-level features were generated using ssGSEA implemented in GSVA v1.50.5 on log_2_(FPKM+1) expression values using the Gaussian-kernel implementation. Only gene sets validated across both cohorts were analysed. The association between each derived feature and TRS was quantified using Kendall's Tau implemented in SciPy v1.11.4. Predictive utility was assessed using mean five-fold cross-validated R^2^. Significance was determined by permutation testing (10,000 label permutations) and BH adjusted. The analysis was repeated using shuffled features to provide a null background.

Features selected from dimensionality reduction via scFoundation and ssGSEA performed best so these were progressed. Two regression approaches were evaluated: a regularised linear ElasticNet model and a non-linear Random Forest model, implemented using scikit-learn (v1.6.1). Model-specific preprocessing pipelines were applied, including feature standardisation for ElasticNet and mutual information-based feature selection for Random Forest, with the number of selected features optimised during hyperparameter tuning. Hyperparameters were optimised within the training cohort (n = 90) using Bayesian optimisation (scikit-optimize v0.10.2) and assessed through repeated five-fold cross-validation (five repeats). The robustness of model selection was further evaluated using nested five-fold cross-validation. Following optimisation, models were retrained on the full training dataset and evaluated on an independent held-out test cohort (n = 30). Model performance was assessed using the coefficient of determination (R²) and root mean square error (RMSE).

**Calculating axis scores**

To enable scoring of individual samples, JBS axis scores were calculated from the dot product of the log_2_-transformed expression values (FPKM) of JBS genes and the PC1 loading vector resulting from applying uncentered and unscaled prcomp (from base R stats v4.3.0) to the JBS gene log2 fold-change gene matrix for the Discovery cohort (n = 96 patients) . These same PC1 loadings were applied to all subsequent bulk, single-cell and spatial transcriptomic datasets to ensure comparability of JBS Axis scores across analyses with a pseudocount of 0.01 for bulk and 1 for cells and Visium spots. This linear transformation preserves the relative ordering of samples along the JBS Axis whilst aligning TPM-derived values more with FPKM-derived scores (R2: 0.97).

Similarly, PC1 loadings for the PN-MES axis were obtained from Wang et al. and samples were projected onto this axis via the dot product of the sample’s bulk RNA-seq gene expression vector and the PC1 loadings, restricted to the intersection of genes present in both datasets.

**Gaussian Mixture Modelling and Polarisation Metrics**

Gaussian mixture models (GMMs) were fitted using the GaussianMixture implementation in scikit-learn (v1.6.1) with default parameters, 20 random initialisations, and a fixed random seed of 11. Model fit was evaluated using the Bayesian Information Criterion (BIC), with lower values indicating improved fit while penalising model complexity. Models with increasing numbers of components were compared, and the change in BIC between one- and two-component models (ΔBIC) was used to assess evidence for bimodality. Distribution polarisation was quantified using complementary metrics including Ashman’s D, which measures separation between Gaussian components, posterior assignment entropy, which captures uncertainty in component membership, and the silhouette coefficient, which evaluates cluster separation and cohesion. Higher Ashman’s D and silhouette values, together with lower entropy, indicate stronger segregation between components. Confidence intervals were estimated by bootstrap resampling (n=200) performed independently for each cohort and tumour stage using a fixed random seed of 11. Mean metric values and standard errors were calculated across bootstrap replicates, and 95% confidence intervals were estimated assuming approximate normality of the bootstrap distributions (mean ± 1.96 × SEM). Differences between primary and recurrent tumours were assessed by comparing confidence interval overlap.

**Compositional Analysis**

Cell type compositions were computed as the proportion of each cell type relative to the total cell count. Similarly for tumour compartment compositions (e.g. isolating

malignant or non-malignant fractions), proportions were calculated relative to the

relevant compartment total. For multivariate regression and correlation analyses,

compositions were transformed using the centred log-ratio transformation (CLR) to address the unit-sum constraint inherent to compositional data following multiplicative replacement of zeros. Ordinary least squares (OLS) regression models were fit using the statsmodels package (v0.14.4) OLS class, with regression coefficients and 95%confidence intervals extracted from the fitted models.

**Treatment-Associated State Dynamics**

JBS Axis scores were computed for individual cells. For each patient and cell type (or cell state within the malignant compartment), the median JBS Axis score was calculated separately for primary and recurrent samples. The change in median score

through treatment was then computed as the difference between recurrent and primary

values. Pearson correlation was used to assess the relationship between this change in

median JBS Axis score and the change in cell type or state composition (CLR-transformed) across patients.

**Brain Organoids**

40,000 ReNcell-VM neural stem cells were seeded per well in a non-adherent 96-well u-bottom plate in ReNcell NSC Maintenance media supplemented with 20ng/mL FGF-2 and 20 ng/mL EGF. After 48hrs, the spheroids were differentiated by replacing the media with ReNcell NSC Maintenance Media without FGF2 and EGF. Spheroids were differentiated for 7 days. Two GBM cell lines, A172 and M059K, stably expressing GFP were cultured in DMEM and DMEM/F-12 media, respectively, containing 10% FCS. In parallel, both lines were seeded (2000 cells/well) in non-adherent 96-well u-bottom plates for 48 hours. To form GLIoma Brain Organoids (GLIBOs), A172 and M059K spheroids (48hrs post seeding) were added to ReNcell-VM spheroids (after maturation for 7 days) in 96 well non-adherent plates in a 1:1 ratio. Plates were left for 48-72 hours to allow formation of GLIBOs, after which light microscopy was used to confirm the GBM spheroids had merged with the VM-ReNcell spheroids. Seven days after assembloid formation, GLIBOs were dissociated into single cells using the Neural Tissue Dissociation Kit (Miltenyi ) and GFP-expressing GBM cells sorted from non-fluorescent ReNcell-VM cells using the Bigfoot Spectral Cell Sorter (Thermo Fisher Scientific). RNA was extracted from GBM cells using the Single Cell RNA Purification Kit (Norgen Biotek) according to the manufacturer’s instructions and converted to cDNA. cDNA was fragmented, converted to libraries using the NEBNext kit and sequenced to 50 million 100bp paired end reads per sample on an Illumna NextSeq 2000.

**Chromatin profiling**

Longitudinal GBM samples were acquired from The Walton Centre and Lancashire Teaching Hospitals National Health Service (NHS) Foundation Trusts via the Brain Tumour Northwest Tissue Banks. Tissue (2-5 mg per reaction) from fresh frozen longitudinal GBMs was minced on ice, fixed in 0.1% formaldehyde for 2 min, and quenched with 125 mM glycine for 5 min at room temperature. Following washing, tissue was mechanically dissociated in wash buffer containing spermidine and protease inhibitors using a Dounce homogenizer. Single-cell suspensions were assessed microscopically and filtered where necessary to remove aggregates. Cells were washed, pelleted, and resuspended in 100 μL wash buffer per reaction before proceeding with the standard CUT&RUN protocol. CUT&RUN was performed by the Epigenomics Profiling Core at MD Anderson Cancer Center following as described(15) with some modifications. Briefly, for each antibody, ~500,000 nuclei isolated from GBM tissues were immobilized on Concanavalin A-coated magnetic beads (Bangs Laboratories) and permeabilized using wash buffer containing 0.02% digitonin (Promega). Bead bound nuclei were incubated with rabbit IgG (Millipore), H3K4me3 (EpiCypher), H3K27me3 (Diagenode) antibodies overnight at 4°C. Next day, targeted chromatin digestion was achieved by pAG-MNase (EpiCypher) binding and incubation with CaCl2 at 4°C followed by CUT&RUN DNA purification using MinElute columns (Qiagen). Libraries from CUT&RUN DNA were prepared using NEBNext® Ultra™ II DNA Library Prep Kit for Illumina® (New England Biolabs) according to the manufacturer's instructions. Sequencing was performed on an Illumina NextSeq 2000 platform to obtain 5-17 million 100nt PE reads per experiment. In addition, H3K4me3 ChIPseq data was acquired via Active Motif (Carlsbad, CA), for the same longitudinal GBM samples from which H3K27me3 profiling was previously acquired (and used again herein), using the same method(1).

**Epigenomic analyses**

Reads were assessed using FastQC, adapter-trimmed with Trim Galore and aligned to the GRCh38 primary assembly genome using Bowtie2. Subsequent filtering retained properly paired, uniquely mapped reads (MAPQ ≥20). PCR duplicates were removed using Picard MarkDuplicates. Processed BAM files were indexed with SAMtools, and alignment statistics were collected using samtools stats, idxstats, and flagstat. Quality metrics from all stages of the workflow were summarised using MultiQC, using and GENCODEv27 gene annotations. Promoter enrichment of histone marks relative to a matched IgG control was assessed using ChromCall, with GRCh38 genome size and the ENCODE hg38 genome blacklist as inputs. Promoters were defined as 2kb windows centred on transcription start site annotated in GENCODEv27. A histone mark was classified as present at a promoter with a positive enrichment score and FDR< 0.2. Correlation of enrichment scores across all promoters was assessed in comparison with high quality ChIPseq GBM data from Weber et al.(16) and outliers were removed.

Figures were generated using matplotlib (v3.6.3), seaborn (v0.13.2) in python, and

ggplot2 (v3.5.1) in R. All statistical analyses were conducted using python, specifically the pingouin package (v0.5.4) and statsannotations (v0.7.2) for statistical computations, or R.
